# The atypical thiol-disulfide exchange protein α-DsbA2 from *Wolbachia pipientis* is a homotrimeric disulfide isomerase

**DOI:** 10.1101/412205

**Authors:** Patricia M. Walden, Premkumar Lakshmanane, Maria Halili, Begoña Heras, Gordon J. King, Andrew E. Whitten, Jennifer L. Martin

**Affiliations:** Institute for Molecular Bioscience, University of Queensland, Brisbane QLD 4072, Australia; School of Biological Sciences, University of Queensland, Brisbane QLD 4072, Australia; Griffith Institute for Drug Discovery, Griffith University, Nathan, QLD 4111, Australia

## Abstract

DiSulfide Bond (DSB) oxidative folding enzymes are master regulators of virulence localized to the periplasm of many Gram-negative bacteria. The archetypal DSB machinery from *Escherichia coli* K12 has a dithiol oxidizing redox relay pair (DsbA/B), a disulfide isomerizing redox relay pair (DsbC/D) and specialist reducing enzymes DsbE and DsbG that also interact with DsbD. By contrast the Gram-negative bacterium *Wolbachia pipientis* encodes just three DSB enzymes. Two of these α-DsbA1 and α-DsbB form a redox relay pair analogous to *E. coli* DsbA/B. The third enzyme α-DsbA2 incorporates a DsbA-like sequence but does not interact with α-DsbB. In comparison with other DsbA enzymes, α-DsbA2 has ∼50 extra N-terminal residues. The crystal structure of α-DsbA2ΔN, the N-terminally truncated form in which these residues are removed confirms the DsbA-like nature of this domain. However, α-DsbA2 does not have DsbA-like activity: it is structurally and functionally different as a consequence of its N-terminal residues. First, α-DsbA2 is a powerful disulfide isomerase and a poor dithiol oxidase – *ie* its role is to shuffle rather than introduce disulfide bonds. Moreover, small-angle X-ray scattering of α-DsbA2 reveals a homotrimeric arrangement. Our results allow us to draw conclusions about the factors required for functionally equivalent enzymatic activity across structurally diverse protein architectures.

## Introduction

Many secreted and outer membrane proteins of prokaryotes rely on disulfide bonds for stability and function (Feige & Hendershot, 2011). The introduction, isomerization and reduction of protein disulfide bonds in bacteria are controlled by <u>d</u>i<u>s</u>ulfide <u>b</u>ond forming (DSB) proteins (Landeta *et al.*, 2018, Inaba, 2009). These DSB proteins are considered master regulators of virulence because they are essential for the folding and activity of diverse virulence factors including bacterial toxins, secretion systems, adhesins, flagella *etc* (Heras *et al.*, 2009).

The classic DSB folding machinery characterised in the model bacterium *Escherichia coli* K-12 comprises two independent periplasmic pathways: the (i) oxidative and (ii) isomerization pathways (Inaba, 2009). In the oxidative pathway, a monomeric thioredoxin (TRX)-fold protein *E. coli* DsbA (EcDsbA) donates its Cys-X-X-Cys active site disulfide bond directly to nascent protein substrates (Zapun *et al.*, 1993, Inaba & Ito, 2002). EcDsbA becomes reduced as a consequence of this reaction and its active site is re-oxidized by a specific interaction with its integral membrane protein partner *E. coli* DsbB (EcDsbB) (Bader *et al.*, 1999).

In the classic isomerization pathway, the *E. coli* homodimeric protein disulfide isomerase DsbC (EcDsbC) reduces and shuffles incorrect disulfide bonds in misfolded proteins to generate correctly folded proteins (Shevchik *et al.*, 1994). Each EcDsbC protomer has a catalytic TRX-fold domain with a characteristic Cys-X-X-Cys active site, and an 87-residue N-terminal region that forms a dimerization domain essential for isomerase activity (McCarthy *et al.*, 2000). EcDsbC forms a redox relay with the integral membrane protein *E. coli* DsbD (EcDsbD) that maintains EcDsbC in its active reduced state (McCarthy *et al.*, 2000). *E. coli* K12 also encodes two specialist reducing enzymes EcDsbG and EcDsbE (Depuydt *et al.*, 2009) that interact with EcDsbD (Missiakas *et al.*, 1995).

Here we focus on one of the DSB proteins encoded by *Wolbachia pipientis w*Mel, a bacterium from the *Rickettsiaceae* family. The *Rickettsiacea*e family are Gram-negative bacteria of the α-Proteobacteria class that establish obligate intracellular infections in arthropods. *W. pipientis* are widespread, found in ∼60% of insect species, and have an extraordinary impact on host biology. Infection results in phenotypic alterations such as cytoplasmic incompatibility, feminization or reduction of lifespan – all of which contribute to the bacterium’s own survival (Hilgenboecker *et al.*, 2008). Several *Wolbachia* strains have been shown to block the transmission of mosquito-borne viruses and are being trialled as biocontrol agents aimed at eradicating vector-borne diseases such as dengue, Zika and Chikungunya (reviewed in (Flores & O’Neill, 2018)).

The *W. pipientis w*Mel strain encodes two DsbA-like proteins, α-DsbA1 and α-DsbA2, and an integral membrane protein α-DsbB (Walden *et al.*, 2013). Unlike *E. coli*, this strain does not encode obvious homologues of DsbC or DsbD (though all other *W. pipientis* strains do encode a DsbD homologue). Of the two encoded DsbAs, α-DsbA1 has been characterized and shown to form a redox relay with α-DsbB (Walden *et al.*, 2013) that resembles the redox relay between EcDsbA and EcDsbB. In contrast, α-DsbA2 – which is highly conserved in *Wolbachia* – does not interact with α-DsbB and it has a long N-terminal region compared with α-DsbA1 (Walden *et al.*, 2013).

The N-terminal region of EcDsbC forms a dimerization domain essential for disulfide isomerase activity. Two other DsbA-like proteins with N-terminal extensions are disulfide isomerases and thought to be dimeric (*Legionella* DsbA2 (Kpadeh *et al.*, 2015), *Caulobacter* ScsC (Cho *et al.*, 2012)). In addition, *P. mirabilis* ScsC is DsbA-like with an N-terminal extension, and has disulfide isomerase activity. However its N-terminal residues interact to form a homotrimer (Furlong *et al.*, 2017). In each of these cases, the N-terminal regions are essential for oligomerisation and protein disulfide isomerase activity. We hypothesised that the N-terminal region of *Wolbachia* α-DsbA2 would also impart disulfide isomerase activity by forming an oligomerisation domain.

Here we report structural and functional studies of *Wolbachia* α-DsbA2. We used two constructs: (1) FL α-DsbA2, the full-length mature α-DsbA2 (lacking the signal peptide that directs the protein to the periplasm) comprising residues 15-252 and (2) α-DsbA2ΔN a truncated form of α-DsbA2 (lacking both the signal sequence and the 50-residue N-terminal region) comprising residues 64-252. Our results show that FL α-DsbA2 is a strong protein disulfide isomerase and that removal of the N-terminal residues eliminates this activity but gives rise to weak dithiol oxidase activity. The crystal structure of truncated α-DsbA2ΔN reveals a classic monomeric DsbA-like architecture. However, SAXS models of FL α-DsbA2 are consistent with a trimer forming through the interaction of the N-terminal residues.

## Results

### The N-terminus of *Wolbachia* α-DsbA2 is predicted to be helical

The 87-residue N-terminal region of the archetypal disulfide isomerase EcDsbC adopts a β-sheet dimerisation domain with a helix that links to the catalytic domain (McCarthy *et al.*, 2000). The presence of a detectable DsbA-like domain and an extended N-terminal region in the *Wolbachia* α-DsbA2 sequence suggested that – like EcDsbC – the 50 residue N-terminal region might act as a dimerisation domain. However, the predicted secondary structure of the α-DsbA2 N-terminal region has no structural relationship with the equivalent region of EcDsbC (**Figure 1A**).

**Figure 1.**
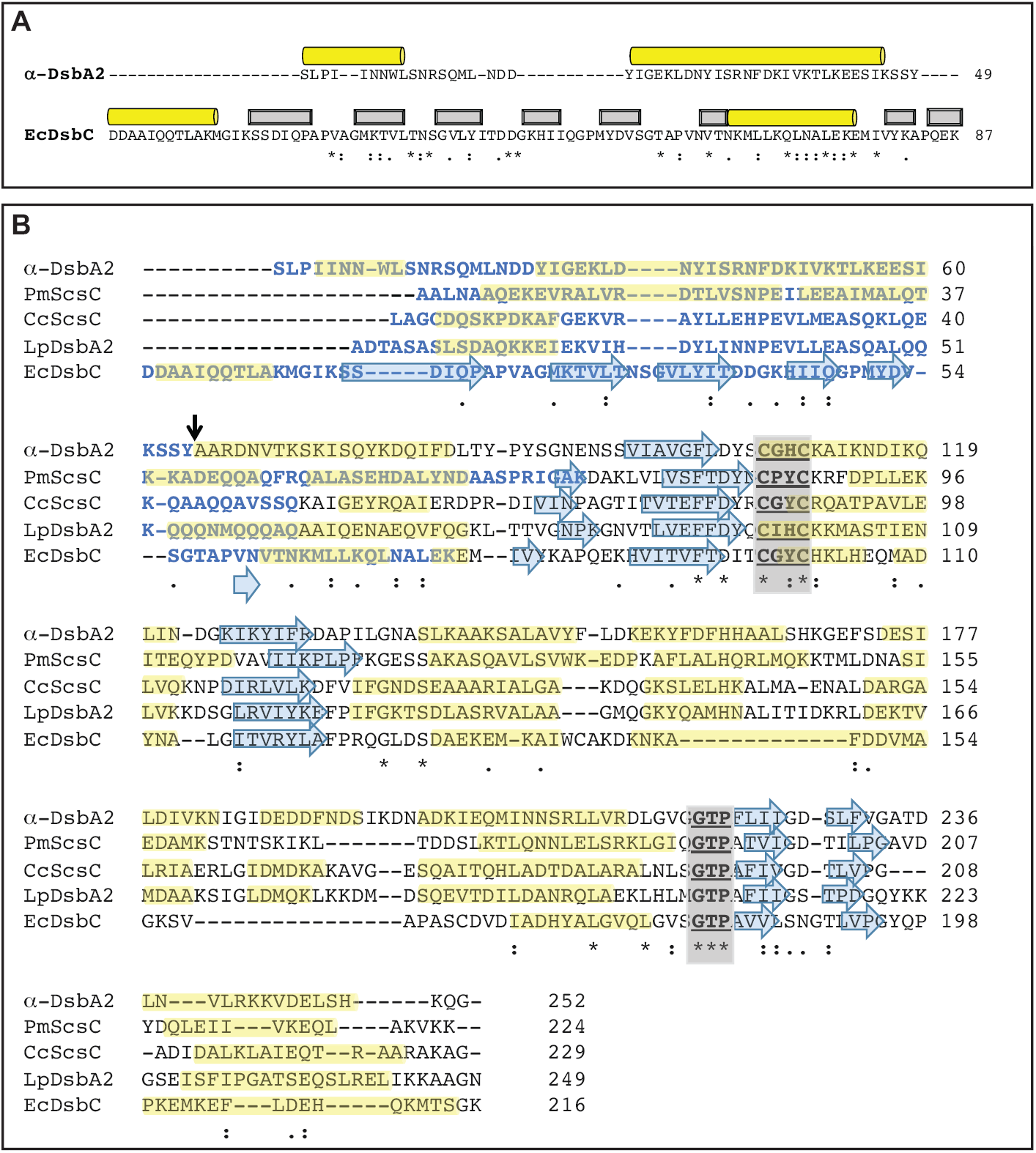
**A) Sequence alignment of the N-terminal residues of α-DsbA2 and EcDsbC.** Sequences were aligned using Clustal Omega (Sievers *et al.*, 2011). Secondary structure was determined from the structure of EcDsbC or from predictions for α-DsbA2 using JPred (Drozdetskiy *et al.*, 2015). α-helices are shown as yellow cylinders and β-strands as grey boxes. Identical residues are marked with an asterisk. **B) Sequence alignment of α-DsbA2, PmScsC, CcScsC, LpDsbA2 and EcDsbC.** Sequences were aligned using Clustal Omega (Sievers *et al.*, 2011). The sequence identities for α-DsbA2 are 21%, 19%, 20%, and 21% with PmScsC, CcScsC and LpDsbA2, and EcDsbC, respectively. The N-terminal residues of each protein sequence are shown as blue bold letters, α-helices are highlighted in yellow and β-strands as blue arrows. The catalytic active site and *cis*-Pro loop residues are bold and underlined. The vertical black arrow indicates the start of the catalytic (TRX) domain in α-DsbA2.

Recently, three other DsbA-like proteins have been shown by gel filtration to be oligomeric as a consequence of their N-terminal residues, and all three are functional disulfide isomerases. These are *Legionella pneumophila* DsbA2 [LpDsbA2, probably dimeric, with a ∼50 residue N-terminal region predicted to be helical (Kpadeh *et al.*, 2015)], *Caulobacter crescentus* ScsC [probably dimeric, ∼60 residue N-terminal region predicted to be helical (Cho *et al.*, 2012)] and *Proteus mirabilis* ScsC [confirmed trimeric protein, ∼60 residue helical N-terminal region (Furlong *et al.*, 2017)]. An alignment of all five proteins (α-DsbA2, LpDsbA2, CcScsC, PmScsC and EcDsbC) is provided in **Figure 1B**.

### α-DsbA2 is redox active and has disulfide isomerase activity as a consequence of its N-terminal residues

We investigated the redox properties of FL α-DsbA2 and truncated α-DsbA2ΔN. The redox potential gives important information about a protein’s propensity to acquire electrons from its substrate, and thereby become reduced. We determined the standard redox potentials of FL α-DsbA2 and α-DsbA2ΔN relative to the redox potential of glutathione (−240 mV) (**Figure 2A)**. From these data, the *K*_eq_ for FL α-DsbA2 was calculated to be 2.16 ± 0.15 × 10^−4^ M corresponding to a redox potential of −131 mV at pH 7.0. The calculated *K*_eq_ for α-DsbA2ΔN was a little more oxidizing, 8.71 ± 0.12 × 10^−5^ corresponding to a redox potential of −122 mV. By comparison, the redox potential for *Wolbachia* α-DsbA1 is more reducing, E^0^ = –163 mV (Kurz *et al.*, 2009). Thus both the FL and truncated α-DsbA2 proteins have redox potentials that are similar to monomeric EcDsbA [E^0^ = –122 mV (Mossner *et al.*, 1998)] and dimeric EcDsbC [(E^0^ = –129 mV (Zapun *et al.*, 1995)].

**Figure 2.**
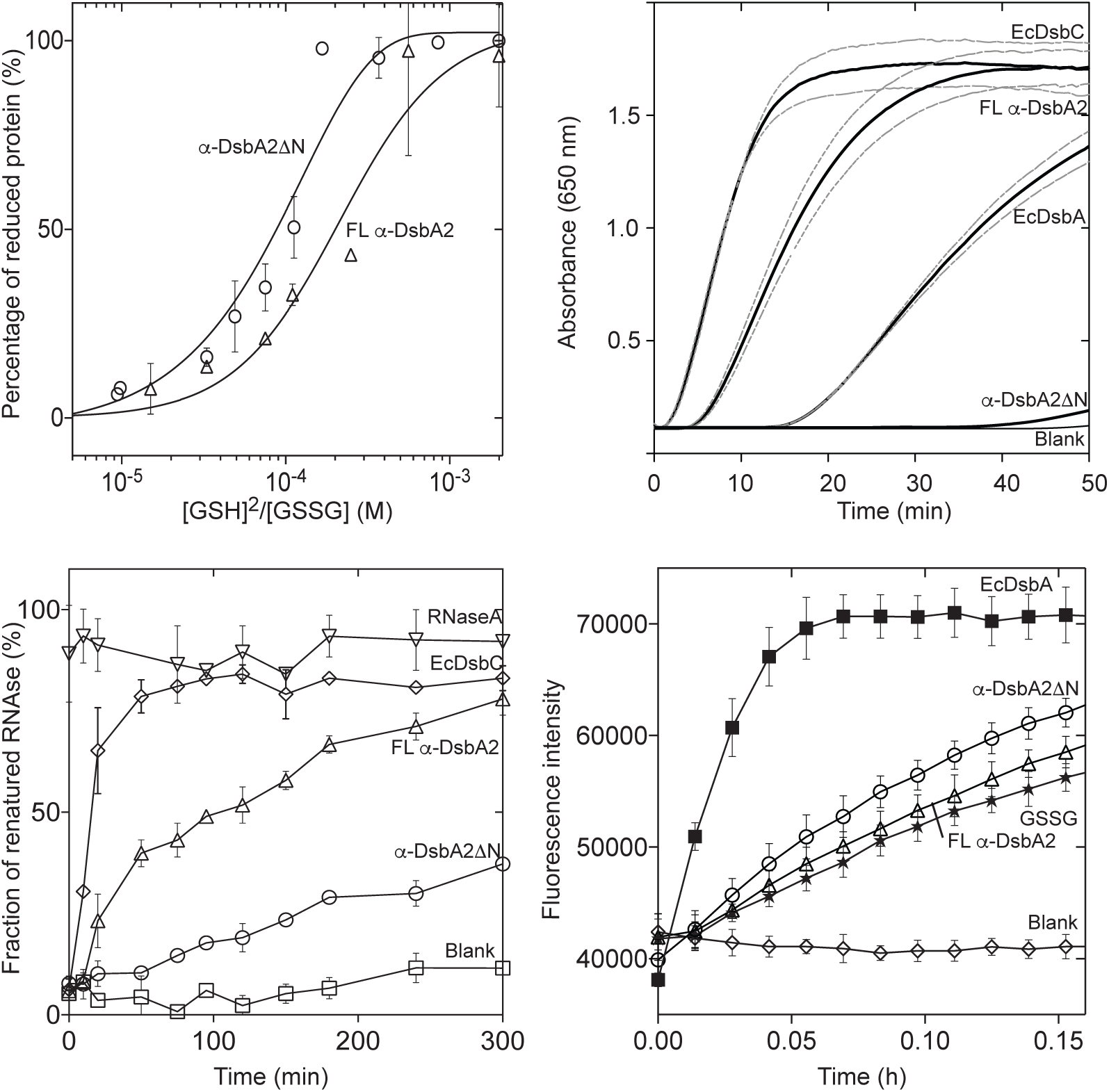
Redox properties of α-DsbA2. **A) Redox potential.** The equilibrium between reduced and oxidized FL α-DsbA2 (triangle) and α-DsbA2ΔN (circle) are shown. Data are presented as the mean ± SD of two measurements. **B) Protein disulfide reductase activity.** The insulin reduction assay was performed to measure the ability of α-DsbA2 variants to reduce insulin. The reduction of insulin catalyzed by the protein or non-catalyzed (blank) was monitored at 650 nm over 50 min. The mean (black line) and standard error (light grey lines) of three replicate measurements are shown. Disulfide oxidase (EcDsbA) and disulfide isomerase (EcDsbC) activities are shown for comparison. **C) Protein disulfide isomerase activity.** Isomerization of scRNase A in the presence of EcDsbC (diamond), FL α-DsbA2 (triangle) or α-DsbA2ΔN (circle), positive control (RNase, inverse triangle) and blank (scRNase without enzyme, square). Data are presented as the mean ± SD of three replicate measurements. **D) Dithiol oxidase activity**. Fluorescence curves showing dithiol oxidation (in the presence of GSSG) of a model peptide by: EcDsbA (positive control, square), α-DsbA2ΔN (circle), FL α-DsbA2 (triangle), GSSG (without enzyme *ie* negative control, star), and α-DsbA2 without peptide (Blank, diamond). Data are presented as the mean ± SD of three replicate measurements.

DSB enzymes are active to varying degrees in the standard disulfide reductase assay. We found that half of the insulin in solution was reduced by FL α-DsbA2 (which is in the reduced form due to DTT in the solution) after ∼15 min (**Figure 2B)**. By comparison, EcDsbC reduced half the insulin within ∼6 min and the oxidase EcDsbA (which is in the reduced form due to DTT in the solution), reduced half of the insulin after ∼35 min. The activity of α-DsbA2ΔN in this assay was negligible, and comparable to the negative control. These data show that FL α-DsbA2 is redox-active and the N-terminal residues are critical for disulfide reductase activity.

We next assessed the protein disulfide isomerase activity of FL α-DsbA2 and α-DsbA2ΔN by following the reactivation of scrambled RNase (scRNase) in an *in vitro* assay. FL α-DsbA2 recovered 73% ± 3% of RNase activity after ∼5 hours, compared to the positive control EcDsbC which recovered 85% ± 5% of RNase activity over the same time period, though reached that level after 100 min and then plateaued (**Figure 2C**). By comparison, truncated α-DsbA2ΔN recovered just 35% ± 3% of RNase activity relative to native refolded RNase over that period, similar to that reported for the oxidase enzyme EcDsbA (Shouldice *et al.*, 2011). Therefore, we conclude that α-DsbA2 is a protein disulfide isomerase and that its disulfide isomerase activity requires the presence of the N-terminal residues.

We also investigated whether FL α-DsbA2 or α-DsbA2ΔN demonstrated dithiol oxidase activity, by measuring their ability to catalyse disulfide bond formation in an *in vitro* assay. Specifically, we measured the ability of FL α-DsbA2 or α-DsbA2ΔN to oxidize the cysteines of a model peptide in the presence of oxidised glutathione (GSSG). Compared to EcDsbA–mediated peptide oxidation, FL α-DsbA2 demonstrated negligible peptide oxidation (comparable to GSSG control) whereas α-DsbA2ΔN had weak activity (**Figure 2D**) under the experimental conditions of this assay.

### The crystal structure of α-DsbA2ΔN reveals a canonical DsbA architecture

We were unable to generate crystals of the full-length mature α-DsbA2. However, crystals of truncated α-DsbA2ΔN did grow and the crystal structure was solved by molecular replacement to a resolution of 2.25 Å (**Table 1, Figure 3A**).

**Table 1.**
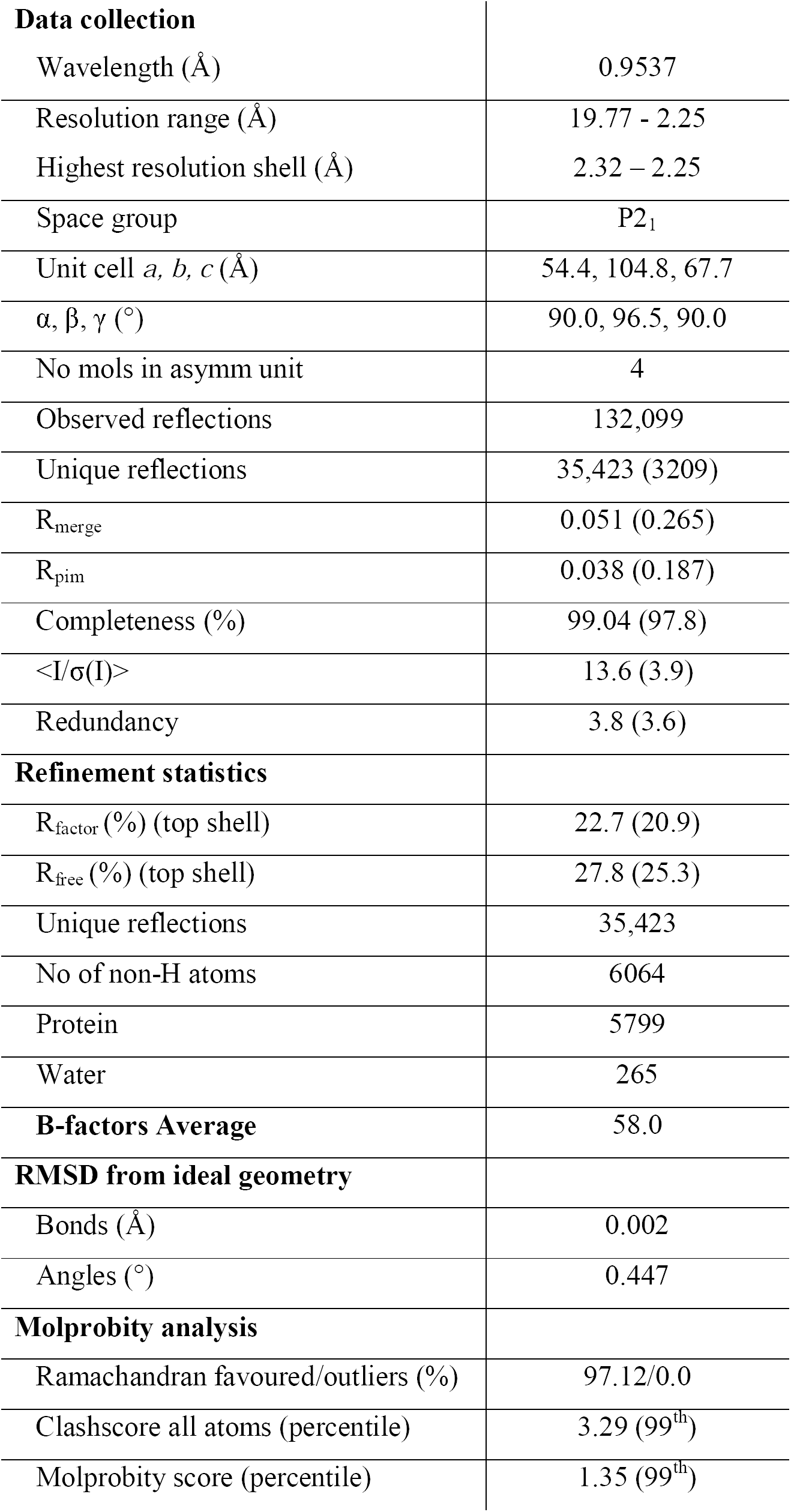
X-ray data collection and refinement statistics for α-DsbA2ΔN

|  |  |
| --- | --- |
| **Data collection** |  |
| Wavelength (Å) | 0.9537 |
| Resolution range (Å) | 19.77 - 2.25 |
| Highest resolution shell (Å) | 2.32 – 2.25 |
| Space group | P2<sub>1</sub> |
| Unit cell *a*, *b*, *c* (Å) | 54.4, 104.8, 67.7 |
| $\alpha$, $\beta$, $\gamma$ (°) | 90.0, 96.5, 90.0 |
| No mols in asymm unit | 4 |
| Observed reflections | 132,099 |
| Unique reflections | 35,423 (3209) |
| R<sub>merge</sub> | 0.051 (0.265) |
| R<sub>pim</sub> | 0.038 (0.187) |
| Completeness (%) | 99.04 (97.8) |
| <I/$\sigma$(I)> | 13.6 (3.9) |
| Redundancy | 3.8 (3.6) |
| **Refinement statistics** |  |
| R<sub>factor</sub> (%) (top shell) | 22.7 (20.9) |
| R<sub>free</sub> (%) (top shell) | 27.8 (25.3) |
| Unique reflections | 35,423 |
| No of non-H atoms | 6064 |
| Protein | 5799 |
| Water | 265 |
| **B-factors Average** | 58.0 |
| **RMSD from ideal geometry** |  |
| Bonds (Å) | 0.002 |
| Angles (°) | 0.447 |
| **Molprobit analysis** |  |
| Ramachandran favoured/outliers (%) | 97.12/0.0 |
| Clashscore all atoms (percentile) | 3.29 (99<sup>th</sup>) |
| Molprobit score (percentile) | 1.35 (99<sup>th</sup>) |

**Figure 3.**
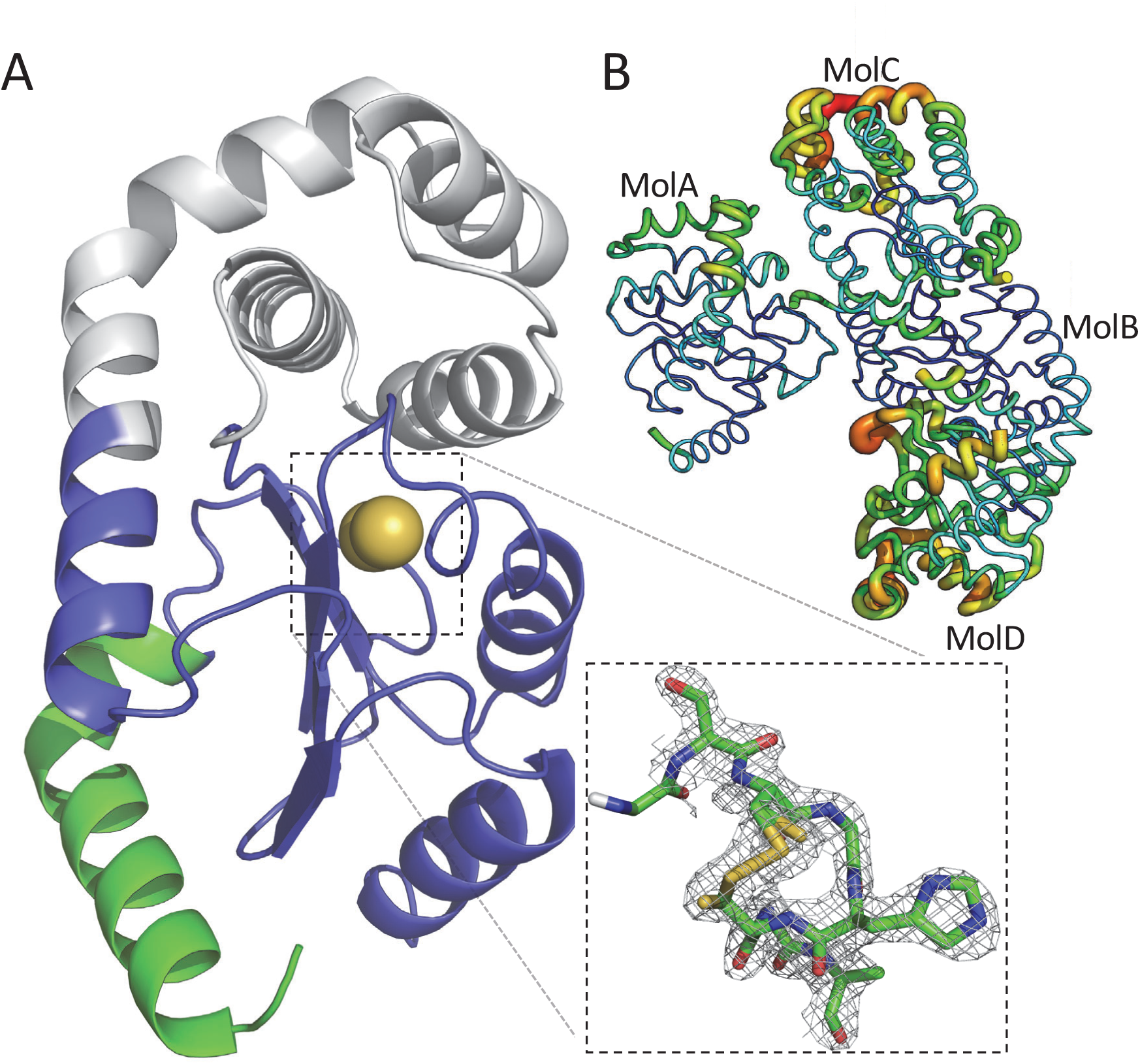
Structure of α-DsbA2ΔN. **A)** Ribbon representation of α-DsbA2ΔN chain B showing the TRX domain in blue and catalytic cysteine sulfur atoms illustrated as yellow spheres. The helical insertion is highlighted in grey, and the N-terminus of the truncated protein is shown in light green. **Dashed lined box:** Active site cysteines Cys107 and Cys110 of chain B were modeled in a mixed redox state (shown for chain A, with the 2F_o_–F_c_ electron density map contoured at 1.0 σ). **B)** The four molecules in the asymmetric unit. The relative crystallographic temperature factors of the refined coordinates are shown by backbone thickness (low-to-high shown as thin-to-thick) and colour (low–to–high coloured from blue–to–red).

The final refined structure of α-DsbA2ΔN has four molecules in the asymmetric unit (chains A, B, C, D) (**Figure 3B**), each of which features the canonical DsbA architecture of a TRX-fold with an inserted 4-helical domain and a connecting helix. As in other DsbA-like crystal structures the Cys-Gly-His-Cys active site is located at the N-terminal end of helix α3 in the TRX domain (present as a mixture of reduced and oxidized forms, **Figure 3A, dashed line box**).

Using this crystal structure as a probe, the highest-scoring DALI match (1 March 2018) was the trimeric protein PmScsC (5IDR, molecule A) with Z score 25.8, RMSD 1.5 Å for 178 C_α_-atoms, and 24% sequence identity. After PmScsC, the next highest DALI hit was an uncharacterised monomeric DsbA-like protein from *Silicibacter pomeroyi* (from the family *Rhodobacteraceae*) (3GYK molecule A) Z score 24.8 RMSD 1.5 Å for 169 C_α_-atoms and 30% sequence identity. The third highest hit was the monomeric protein SeScsC (4GXZ, molecule C) Z score 24.1 RMSD 1.6 Å for 166 C_α_-atoms, and 25% sequence identity (this was used as the molecular replacement model to solve the α-DsbA2ΔN crystal structure).

By comparison, superimposition of the α-DsbA2ΔN structure onto the archetypal disulfide isomerase EcDsbC (1EEJ, molecule A) using DALI gave a Z score of 13.5 and RMSD of 3.3 Å for 147 C_α_-atoms (Maiti *et al.*, 2004). These data indicate that these two disulfide isomerases are structurally very different.

### Small angle X-ray scattering shows α-DsbA2 is a homotrimer as a consequence of its N-terminal region

Although we were unable to crystallise the full-length protein, we were able to obtain low-resolution structural information using small-angle X-ray scattering (SAXS) data from both FL α-DsbA2 and α-DsbA2ΔN (**Table 2**). Guinier plots (**Figure 4A, inset**) reveal a linear trend, consistent with both samples being monodisperse.

**Table 2.** SAXS data collection and analysis details

| Data Collection Parameters | α-DsbA2ΔN | α-DsbA2 |
| --- | --- | --- |
| Instrument | SAXS-WAXS (Australian Synchrotron) |  |
| Beam geometry | Point |  |
| Wavelength (Å) | 1.033 | 1.127 |
| Sample to detector distance (m) | 1.575 | 1.575 |
| $q$-range (Å<sup>-1</sup>) | 0.01 – 0.57 | 0.01-0.48 |
| Exposure time (s) | 16 (8 × 2 s exposures) | 40 (8 × 5 s exposures) |
| Configuration | Concentration series from<br>96-well plate | SEC-SAXS<br>Column: WTC-030S5<br>Flow: 0.5 mL/min<br>Inject: 100 µL,<br>19.8 mg/mL |
| Protein concentration range (mg/mL) | 1.25 – 5.00 | 0.35 – 2.20 |
| Temperature (K) | 283 | 293 |
| Absolute intensity calibration | Water |  |
| Sample details |  |  |
| Extinction coefficient ($A_{280}$, 0.1%<br>$w/v$) | 0.494 | 0.762 |
| Partial specific volume (cm<sup>3</sup>g<sup>-1</sup>) | 0.735 | 0.735 |
| Contrast, $\Delta\rho$ (10<sup>10</sup> cm<sup>-2</sup>) | 2.88 | 2.89 |
| Molecular mass [from sequence]<br>(kDa) | 21.1 | 27.1 |
| Protein concentration (mg/mL)* | 1.25 | 0.75 |
| Structural parameters |  |  |
| $I(0)$ (cm<sup>-1</sup>) [from Guinier] | 0.0215 ± 0.0001 | 0.04846 ± 0.00003 |
| $R_g$ (Å) [from Guinier] | 18.7 ± 0.1 | 28.0 ± 0.1 |
| $I(0)$ (cm<sup>-1</sup>) [from $p(r)$] | 0.0215 ± 0.0001 | 0.04845 ± 0.00002 |
| $R_g$ (Å) [from $p(r)$] | 18.8 ± 0.1 | 27.9 ± 0.1 |
| $D_{max}$ (Å) | 63 ± 3 | 87 ± 3 |
| Porod volume (Å<sup>3</sup>) | 27500 ± 1500 | 91300 ± 4500 |
| Molecular mass determination |  |  |
| Molecular mass [from $I(0)$] (kDa) | 23 ± 1 | 87 ± 5 |
| Molecular mass [from Porod] (kDa) | 23 ± 1 | 75 ± 5 |

\* α-DsbA2 Δ65: $I(0) = 0.0427 \pm 0.0001 \text{ cm}^{-1}$ , $R_g = 18.9 \pm 0.1 \text{ Å}$ , $M = 23 \text{ kDa}$ (2.5 mg/mL); $I(0) =$ $0.0890 \pm 0.0002 \text{ cm}^{-1}$ , $R_g = 19.2 \pm 0.1 \text{ Å}$ , $M = 24 \pm 2 \text{ kDa}$ (5.0 mg/mL); There is a small but significant systematic change in $R_g$ and $M$ in the concentration range measured, but based on the trend, 1.25 mg/mL was deemed to be free of concentration dependent effects; FL $\alpha$ DsbA2: At the peak concentration of ~2.2 mg/mL, $R_g = \sim 27.6 \text{ Å}$ , which increases and plateaus at $R_g = \sim 28.0 \text{ Å}$ at a concentration of between 0.35 – 1.15 mg/mL. This range was deemed to be free of concentration dependent effects and all eight frames collected over these concentrations were combined, where the average concentration was 0.75 mg/mL.

**Figure 4.**
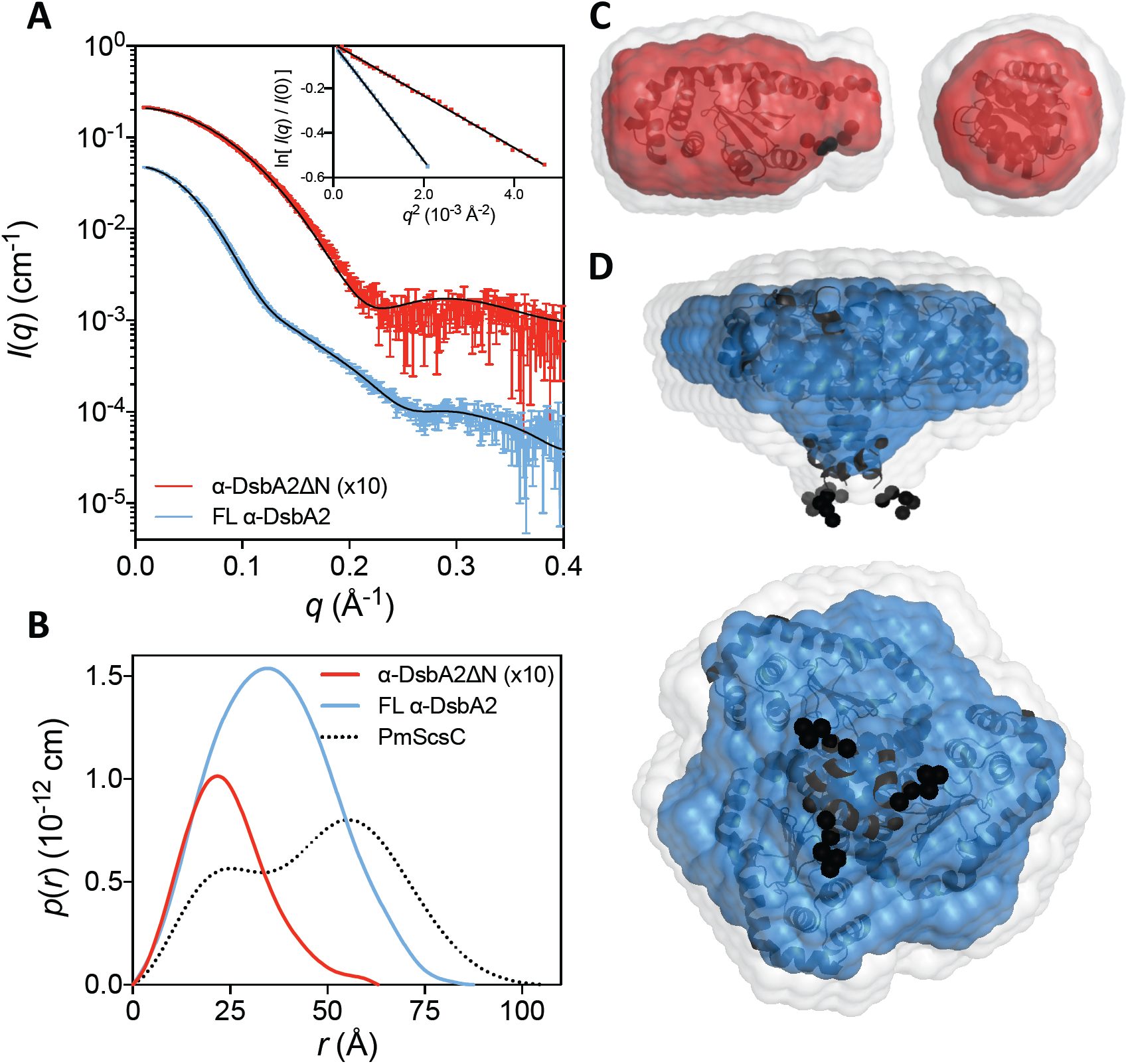
Small angle X-ray scattering data for α-DsbA2ΔN and FL α-DsbA2. **A)** Measured scattering data for α-DsbA2ΔN (red, multiplied by a factor of 10 for clarity) and FL α-DsbA2 (blue). The scattering profile of rigid-body models are shown as solid black lines overlaid on the scattering data for α-DsbA2ΔN (χ^2^ = 3.88; CorMap test (Franke *et al.*, 2015), 294 points, *C*=66, *P*=0.000) and FL α-DsbA2 (χ^2^ = 1.37; CorMap test, 320 points, *C*=13, *P*=0.037). **Inset:** Guinier plot for α-DsbA2ΔN (red, *R*^2^ = 0.999) and FL α-DsbA2 (blue, *R*^2^ = 1.000). **B)** The pair-distance distribution function, *p*(*r*), derived from the scattering data are indicative of a globular structure with a maximum dimension of ∼63 Å for α-DsbA2ΔN (red) and ∼87 Å for FL α-DsbA2 (blue). For reference, the experimental *p*(*r*) for the trimeric PmScsC is also shown (dotted line). **C)** Probable shape of α-DsbA2ΔN (monomeric) obtained from the filtered average of 16 dummy-atom models (red envelope): χ^2^ = 1.038 ± 0.002; NSD = 0.446 ± 0.021; Resolution = 17 ± 2. **D)** Probable shape of FL α-DsbA2 obtained from the filtered average of 9 dummy-atom models (blue envelope): χ = 1.202 ± 0.003; NSD = 0.602 ± 0.027; Resolution = 30 ± 2. Images in **C** and **D** were generated using PyMol, where the grey shapes represent the total volume encompassed by the aligned dummy-atom models and the corresponding rigid-body model is shown aligned to the filtered model (flexible regions are represented by chains of black spheres).

Bacterial disulfide isomerases characterised to date are reported to be either dimeric (EcDsbC, CcScsC, LpDsbA2) or trimeric (PmScsC). We were therefore interested to determine the molecular mass of the full-length protein and identify whether it is dimeric or trimeric. Using SAXS data, the molecular mass of truncated α-DsbA2ΔN estimated to be ∼23 kDa from *I*(0) (Orthaber *et al.*, 2000) and the Porod volume (Fischer *et al.*, 2010), which is very close to the expected mass for an α-DsbA2ΔN monomer (21 kDa) and consistent with the crystal structure we report here. However, the molecular mass of FL α-DsbA2 estimated from *I*(0) and the Porod volume (75 kDa and 87 kDa respectively) are consistent with the mass of a homotrimer (81 kDa) rather than a homodimer (54 kDa).

The *p*(*r*) for FL α-DsbA2 shows a single peak with a maximum dimension of ∼87 Å (**Figure 4B**) whereas α-DsbA2ΔN demonstrates a single peak with a significantly smaller maximum dimension of 63 Å. The larger dimension and shifting of the position of the peak in the *p*(*r*) for FL α-DsbA2 are consistent with the formation of a higher-order oligomer. While these data indicate the FL α-DsbA2 oligomer is trimeric, this homotrimer differs from that of PmScsC (Furlong *et al.*, 2017), which has a bimodal pair-distance distribution function (**Figure 4B, dotted line**).

Both dummy atom and rigid body modelling were used to determine low-resolution solution structures of FL α-DsbA2 and α-DsbA2ΔN. The averaged and filtered dummy-atom model of α-DsbA2ΔN shows very good agreement with the α-DsbA2ΔN rigid-body model (composed of the crystal structure plus missing residues at the N- and C-termini, **Figure 4C**). A comparison of the scattering data with the rigid-body model scattering profile (**Figure 4A**, red curve and black line) shows good correspondence, though with a small systematic difference between the two curves that could be indicative of a low level of sample impurity or possibly a small difference between the solution and crystal structures.

The averaged and filtered dummy-atom model of FL α-DsbA2 reveals a disk-like structure with a small protrusion at the centre. Alignment of this model with the FL α-DsbA2 rigid-body model (**Figure 4D**) shows excellent correspondence; the N-terminal oligomerisation domain of the rigid-body model coincides with the protrusion in the dummy-atom model, and the catalytic domains are positioned around the main disk. The scattering data for FL α-DsbA2 shows excellent correspondence with the rigid-body model scattering profile (**Figure 4A**, blue curve and black line).

The two homotrimeric disulfide isomerase enzymes *Wolbachia* FL α-DsbA2 and PmScsC nevertheless have distinct solution structures. FL α-DsbA2 is disc-like with the three protomers tightly arranged into a compact shape (**Figure 5A, B**), whereas PmScsC has a more open arrangement (**Figure 5C, D)** (Furlong *et al.*, 2017).

**Figure 5.**
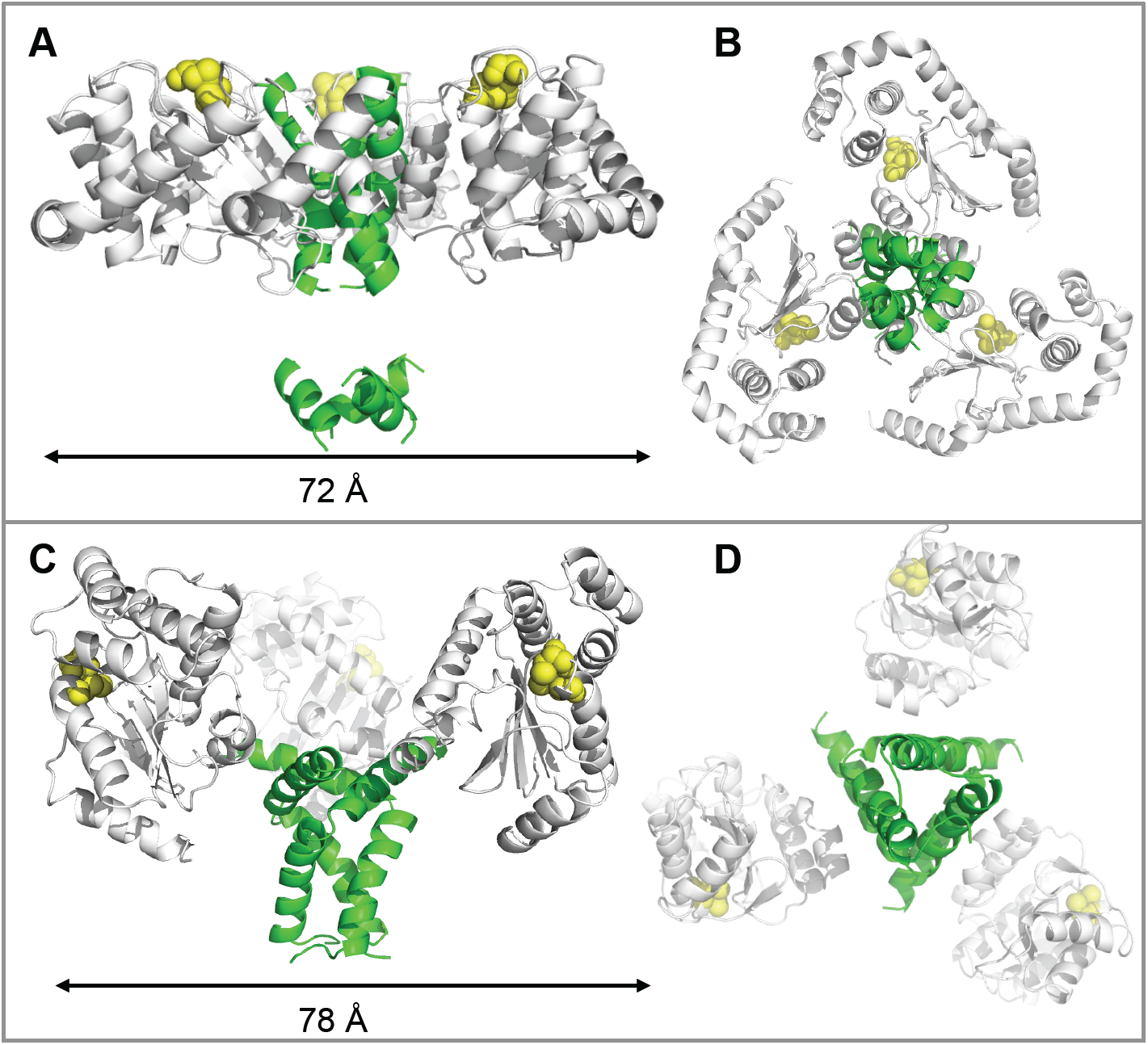
Comparison of rigid body models of FL α-DsbA2 and PmScsC. Model of FL α-DsbA2 from **A)** the side and **B)** bottom view. Model of PmScsC (SASDBD code: SASDB94) from **C)** side and **D)** bottom view. For both FL α-DsbA2 and PmScsC, the N-terminal trimerisation domain is shown as a green ribbon, the catalytic domain as a white ribbon, and active site residues as yellow spheres. The regions treated as flexible linkers in the rigid-body model are not shown, which is the reason for the apparent gap in the N-terminal trimerisation domain shown in panel A.

## Discussion

The *w*Mel strain of *W. pipientis* encodes two DsbA-like proteins. One of these, α-DsbA1 has been shown previously to be functionally similar to EcDsbA – it catalyses disulfide formation and forms a redox pair with a membrane protein partner α-DsbB (Walden *et al.*, 2013). Despite sequence similarity, the second protein α-DsbA2, is not DsbA-like. It does not catalyse disulfide formation in the standard assay (this work) and it does not interact with α-DsbB (Walden *et al.*, 2013).

The unusually long N-terminus of *W. pipientis w*Mel α-DsbA2 is conserved in DsbA2s encoded in bacterial species from the family *Rickettsiaceae*, class α-Proteobacteria. This includes other members from the same class including *Ehrlichia* and *Anaplasma* that live in hosts such as ticks and cause diseases in animals and humans (Wormser *et al.*, 2006). These organisms all encode α-DsbA1 and α-DsbA2 enzymes similar to those encoded in *Wolbachia*. The high degree of conservation of the N-terminal residues in α-DsbA2 suggests that this region has an important function. Here we showed that these N-terminal residues confer trimerisation and disulfide isomerase properties to *W. pipientis w*Mel α-DsbA2.

The enzymatic profile of *W. pipientis w*Mel α-DsbA2 overlaps with that of EcDsbC. Indeed, it may replace EcDsbC functionally since *Wolbachia* strains do not encode a DsbC. Most *Wolbachia* strains encode an EcDsbD-like protein that could act as a redox partner for α-DsbA2. Curiously, a DsbD homologue is not present in the genome of the specific *w*Mel *Wolbachia* strain we investigated. We cannot explain why this might be or what other protein might serve as a reducing partner in this organism.

Other organisms encode a disulfide isomerase like that of *Wolbachia* α-DsbA2 rather than EcDsbC. For example, *Legionella pneumophilia* does not contain a DsbC homologue (Kpadeh *et al.*, 2013, Kpadeh *et al.*, 2015) though like *Wolbachia* it has two DsbA-like proteins, one of which, LpDsbA2, has disulfide isomerase activity. LpDsbA2 has a predicted helical N-terminal extension (**Figure 1B**), and is essential for assembly of the Type 4b Dot/Icm secretion system (Kpadeh *et al.*, 2013, Kpadeh *et al.*, 2015).

We have shown that *W. pipientis w*Mel α-DsbA2 is homotrimeric and has disulfide isomerase activity. Although this is the second example of a homotrimeric TRX-fold disulfide isomerase – the other being PmScsC (Furlong *et al.*, 2017) - these two enzymes are quite distinct. First, α-DsbA2 is structurally different from PmScsC in solution. PmScsC (Furlong *et al.*, 2017) exhibits a bimodal pair-distance distribution function whereas FL α-DsbA2 has a single peak indicating a more globular and compact shape. Both dummy atom and rigid body modelling reveal that FL α-DsbA2 is disc-like with no evidence of the flexibility observed for PmScsC (Furlong *et al.*, 2017). Moreover, PmScsC is part of a highly conserved 4-gene *scs* cluster associated with bacterial copper resistance, including its redox partner PmScsB (Furlong *et al.*, 2018) whereas α-DsbA2 is not part of a gene cluster. Finally, PmScsC is encoded in organisms that also encode DsbC-like enzymes, whereas α-DsbA2 is not. Presumably, PmScsC may play a specific role – perhaps it has a specific substrate associated with copper sensitivity – whereas α-DsbA2 may not.

Although these three structurally characterised disulfide isomerases - EcDsbC, PmScsC and α-DsbA2 - differ considerably in their structures, we can draw some broad conclusions about the factors that contribute to their functionally equivalent enzymatic activity. The present work supports the notion that strong disulfide isomerase activity requires the presence of at least two catalytic domains in the enzyme. The way these domains are brought together can vary (dimer/trimer; sheet/helix) and there is some limited variation in the catalytic active site motif: CGYC in DsbC and PmScsC; CPYC in CcScsC; CIHC in LpDsbA2; and CGHC in *Wolbachia* α-DsbA2. Thus, Gly or Pro predominate in the Cys+1 position, and Tyr or His predominate in the Cys+2 position. However, this catalytic motif sequence overlaps with that of monomeric dithiol oxidase DsbAs (eg EcDsbA CPHC). The second motif that is highly conserved in TRX-like proteins is the *cis*-Pro motif. Surprisingly, in all five of these disulfide isomerase enzymes the sequence motif is the same GT*c*P. Monomeric DsbA-like oxidases tend to have more variation, with the Gly being highly variable and the Thr often replaced with Val. The most telling sequence feature that discriminates between oxidase and isomerase activity seems to be the addition of 50 or more residues at the N-terminus of the TRX fold that can form an oligomerisation domain.

## Acknowledgements

Structural data reported in this paper were measured on the MX2 and SAXS/WAXS beamlines at the Australian Synchrotron. We thank the beamline staff for their support. We also thank the University of Queensland Remote Operation Crystallization and X-ray (UQ ROCX) Diffraction Facility for access to crystallization facilities. Additionally, we acknowledge Dr Lachlan Casey for SAXS data measurement, Dr Iñaki Iturbe-Ormaetxe for providing *Wolbachia w*Mel DNA and Dr Mareike Kurz for cloning α-DsbA2 into the original expression vector. This work was supported by an Australian International Postgraduate Research Scholarship to PW, and an ARC Australian Laureate Fellowship (FL0992138) to JLM.

## Materials and Methods

### Sequence analysis of α-DsbA2

Primary and secondary structure information was obtained from ExPASy (Swiss Institute of Bioinformatics, Switzerland) using the proteomics server ProtParam (Bairoch *et al.*, 2005) and the server PredictProtein (Rost *et al.*, 2004). Sequences of α-DsbA2, EcDsbC, PmScsC, CcScsC and LpDsbA2 were aligned in ClustalOmega (Thompson *et al.*, 1994). Accession codes: EcDsbC P0AEG6; PmScsC B4EV21; CcScsC Q9A747; α-DsbA2 Q73FL6; LpDsbA2 Q5WVK9.

### Protein expression and purification

FL α-DsbA2 (locus WD1312; GenBank #AE017196) was amplified from *w*Mel genomic DNA by PCR using primers Forward – 5’ TAC TTC CAA TCC AAT GCG ATG AGC TTG CCG ATA ATA 3’ and Reverse – 5’ TTA TCC ACT TCC AAT G CT AGC CTT GCT TGT GAC TTA A 3’ which incorporate overhangs for ligation independent cloning. Truncated α-DsbA2ΔN was amplified using the primers Forward – 5’ TAC TTC CAA TCC AAT GCG GCT CGA GAT AAT GTA ACC 3’ and Reverse – 5’ TTA TCC ACT TCC AAT GCT AGC CTT GCT TGT GAC TTA A 3’. The full-length mature protein without the signal peptide and α-DsbA2ΔN were cloned into the LIC vector pMCSG7 that incorporates a HIS_6_-tag, a linker region containing 8 amino acids, and a TEV-cleavage site at the N-terminus of the inserted gene. The construct was transformed into the expression strain BL21(DE3)pLysS (Life Technologies, USA) to enable over-expression of α-DsbA2 using auto induction in ZYP-5052 media at 30°C (Studier, 2005). Cells were collected using an Avanti J-25I centrifuge (Beckman Coulter, Australia) at 12,000 × g at 4°C for 10 min and frozen at −80°C. α-DsbA2 variants were expressed and purified as described in (Kurz *et al.*, 2009) with minor variation in the lysis, wash and elution buffer. The lysis buffer contains 25 mM Tris-HCl pH 7.5 and 150 mM NaCl plus 25 mM imidazole for the washing buffer, and 250 mM imidazole was included for protein elution.

### Protein disulfide reductase assay

The ability of FL α-DsbA2 and α-DsbA2ΔN to catalyse the reduction of insulin in the presence of DTT was measured *in vitro* (Holmgren, 1979). Insulin comprises two chains, A and B, which are linked via two disulfide bonds. Upon reduction of the disulfide bonds by a high reductase-active catalyst, chain B becomes insoluble and precipitates. FL α-DsbA2, α-DsbA2ΔN, EcDsbC (positive control) or EcDsbA (negative control) were diluted to a final concentration of 10 μM in buffer containing 100 mM NaPO_4_ pH 7.0, 1 mM EDTA and 0.33 mM DTT. Insulin (at a final concentration of 0.13 mM) was added to the cuvette immediately before measurements were taken, and the extent of insulin reduction was monitored by measuring the optical density at 650 nm for 50 min. Three experimental replicates were measured using the same batch of protein and the presented data [mean ± standard deviation error (SD)] is from three measurements.

### Redox potential measurement

2 µM of oxidized FL α-DsbA2 and α-DsbA2ΔN was incubated in fully degassed buffers containing 100 mM NaPO_4_ pH 7.0, 1 mM EDTA and 1 mM glutathione oxidized (GSSG) (Sigma Aldrich, USA) and different concentrations of reduced glutathione (GSH) (20 µM – 5 mM) for 24 h at RT. After incubation, the reactions were stopped with 10% trichloroacetic acid (TCA) (Sigma Aldrich, USA) and the precipitated protein pellets were collected by centrifugation at 16000 x g for 10 min at 4 °C. The pellets were washed with cold acetone and dissolved in buffer containing 50 mM Tris-HCl pH 7.0, 1% SDS and 4 mM 4-acetamide-4’-maleimidylstilbene-2-2’-disulfonate (AMS) (Molecular Probes, USA). The reduced and oxidized forms were separated on a 12% SDS Bis-Tris PAGE (Invitrogen, Australia). The fraction of reduced protein, *R*, was determined from a scanned image of the stained gel using ImageJ (Abramoff, 2004). The equilibrium constant K_*eq*_ was calculated via *R* = ([GSH]^2^/[GSSH])/(*K*_eq_ + ([GSH]^2^/[GSSH])), and the redox potential was calculated via the Nernst equation E^0^ = E^0^_GSH/GSSG_ – (*RT*/*nF*)×ln*K*_eq_, where E^0^_GSH/GSSG_ = −240 mV 240 mV, *R* = 8.314 J K^−1^mol^−1^, *T* = 298 K, *n* = 2 and *F* = 9.649 × 10^4^ C mol^−1^ (Kurz *et al.*, 2009). Three experimental replicates were measured using the different batch of protein and data presented are the mean ± SD from duplicate measurements.

### Dithiol oxidase activity assay

Assays were run on a Synergy H1 Multimode plate reader (BioTek, USA) with excitation at 340nm and emission at 615nm. For time-resolved fluorescence, a 100 µsec delay before reading and 200 msec reading time was employed. The assay was performed in a white 384-well plate (Perkin Elmer). A 25 μL solution containing 3.2 μM EcDsbA and 2 mM GSSG (positive control), or 3 µM FL α-DsbA2 or α-DsbA2ΔN and 2 mM GSSG in 50 mM MES, 50 mM NaCl, 2 mM EDTA pH 5.5 was added to the wells. The assay was initiated by the addition of 25 μL 16 μM peptide (in 50 mM MES, 50 mM NaCl, 2 mM EDTA, pH 5.5) to each well. Measurements were carried out in triplicate using 3 different protein batches and data shown are the mean ± SD error from these three measurements.

### Protein disulfide isomerase assay

The scrambled RNaseA assay was used to detect isomerase activity of FL α-DsbA2, α-DsbA2ΔN or EcDsbC by monitoring the refolding of scrambled RNaseA (Hillson *et al.*, 1984). When the four randomly oxidized disulfides are correctly paired, the RNase is natively folded and active and can convert cyclic cytidine-3,5’-monophosphate (cCMP) into 3’CMP, which can be monitored colorimetrically. In these experiments FL α-DsbA2, α-DsbA2ΔN or EcDsbC (10 µM final concentration) were used in a buffer containing 100 mM sodium phosphate, 1 mM EDTA, pH 7.0, 10 µM dithiothreitol (DTT) and 40 μM of scrambled RNaseA. Scrambled RNaseA was produced as previously described (Kurz *et al.*, 2009). At various time points, 50 µL of the reaction was mixed with 150 µL of cytidine 3’,5’-cyclic monophosphate (3 mM) and hydrolysis activity was monitored using a Biotek H1 plate reader (Millennium Science, USA) at 296 nm and 298 K. Native RNaseA and scrambled RNaseA samples without added enzyme served as positive and negative controls, respectively. Measurements were performed in triplicate using three different batches of protein and presented data is the mean ± SD from three measurements.

### Crystallization and crystal structure determination of α-DsbAΔ65

Crystal screening and optimisation of α-DsbAΔ65 crystals was performed at the UQ ROCX facility (University of Queensland, Australia). For crystallisation, the hanging drop vapour diffusion method was used. Crystals of α-DsbAΔ65 were grown by mixing 1 µL of protein at 20 mg/mL and 1 µL crystallant solution, containing 100 mM Tris pH 7.5, 200 mM NaCl and 20% *w/v* PEG 3350. X-ray data were recorded on the microcrystallography beamline MX2 at the Australian Synchrotron using the Blue-Ice software (McPhillips *et al.*, 2002). Reflections were processed in XDS (Kabsch, 2010), analysed and converted to MTZ in SCALA (Evans, 2006). Phases were obtained by molecular replacement using remote sequence homologues identified by Fold and Function Assignment (FFAS) (Jaroszewski *et al.*, 2011). The top FFAS hit, *Salmonella enterica serovar Typhimurium* ScsC (SeScsC, PDB ID: 4GXZ (Shepherd *et al.*, 2013)) shares 23% sequence identity to α-DsbAΔ65. An initial search using the complete SeScsC PDB coordinates as a model was unsuccessful. Instead a trimmed poly-Ser template that retained the side-chains of residues that were conserved in an alignment of SeScsC with α-DsbA2ΔN was used to phase the structure using PHASER (McCoy *et al.*, 2007) for α-DsbA2ΔN. The root mean square deviation between these two structures is 1.8 Å for 124 equivalent C^α^ atoms. Further refinement was performed using PHENIX (Adams *et al.*, 2010) (Adams *et al.*, 2011) and COOT. Molecular figures were generated in PyMOL (*The PyMOL Molecular Graphics System, Version 1.2r3pre, Schrödinger, LLC.*, LLC. #211). Electrostatic potential figures were generated using APBS (Dolinsky *et al.*, 2004).

RMSD calculations and structural alignments were conducted using PyMOL as well as DaliLite {Holm, 2008 #39}. The data collection and refinement statistics are given in (**Table 1**). The refined model of α-DsbA2ΔN has four molecules in the asymmetric unit with chains A, B, C and D refined with 185, 189, 180 and 180 residues, respectively. Molecules A and B are both well defined in the electron density, but poor density regions in molecules C (residues 188-197) and molecule D (residues 92-98 and 193-197) were not able to improve during refinement with only the Ca atoms and occupancies set to 0 for any other atoms. The different quality of these molecules is evident from the average B-factors of A/B (46 and 41 Å^2^ respectively), and molecules C/D (63 and 70 Å^2^ respectively), (see **Figure 3B**). The catalytic disulfide bonds in molecules A and molecule B were modeled in a mixed redox state (**Figure 3**), whereas in molecules C/D these are modelled as reduced.

### Small angle X-ray scattering of FL α-DsbA2 and α-DsbA2Δ65

SAXS data for α-DsbA2ΔN and FL α-DsbA2 were collected on the SAXS-WAXS beamline at the Australian Synchrotron (Kirby *et al.*, 2013). Data reduction was carried out using Scatterbrain software (v 2.71) (Australian Synchrotron), and corrected for solvent scattering, sample transmission, and detector sensitivity. For α-DsbA2ΔN, serial dilutions of a ∼5 mg/mL stock were loaded into a 96-well plate, while FL α-DsbA2 was measured using an in-line SEC-SAXS set-up (see **Table 2**). The estimated molecular mass was calculated using contrast and partial specific volumes determined from the protein sequences (Whitten *et al.*, 2008). Data processing and Guinier analysis was performed using Primus (v 3.2) (Konarev *et al.*, 2003). The pair-distance distribution function (*p*(*r*)) was generated from the experimental data using *GNOM* (v 4.6) (Svergun, 1992), from which *I*(0), *R*_g_ and *D*_max_ were determined. The program *DAMMIN* (v 5.3) (Svergun, 1999) was used to generate 16 dummy-atom models for each protein, (assuming *C*_1_ symmetry for α-DsbA2ΔN, and *C*_3_ symmetry for α-DsbA2ΔN), which were averaged using the program *DAMAVER* (v 2.8.0) (Volkov & Svergun, 2003), and the resolution of the averaged structures were estimated using SASRES (Tuukkanen *et al.*, 2016). All 16 dummy-atom models were used in the averaging procedure for α-DsbA2ΔN, but only 9 (oblate) of the 16 dummy-atom models were averaged for FL α-DsbA2. Rigid-body modelling was carried out using the program CORAL (v 1.1) (Petoukhov *et al.*, 2012). For α-DsbA2ΔN, residues D71 – L250 were taken from the crystal structure and treated as a rigid unit, while 5 additional residues were included at the N- and C-termini and treated as flexible linkers. For FL α-DsbA2, *C*_3_ symmetry was assumed and L17 – R27 (a model helical segment), D34 – E57 (a model helical segment) and A65 – L250 (from the α-DsbA2ΔN crystal structure) were taken as rigid subunits, while 5 additional residues were added at the N- and C-termini plus the intervening regions between rigid segments and treated as flexible linkers. As oligomerisation occurs through the N-terminal region, and the model helices were amphipathic, L17, I20, W23, I36, L40, I44, F48, V52 and L55 were restrained to be less than 15 Å from the same residue in a symmetry related chain.

